# Arbidol inhibits esophageal squamous cell carcinoma growth *in vitro* and *in vivo* through suppressing ataxia telangiectasia and Rad3-related protein kinase

**DOI:** 10.1101/2022.02.15.480509

**Authors:** Ning Yang, Xuebo Lu, Yanan Jiang, Lili Zhao, Donghao Wang, Yaxing Wei, Yin Yu, Myoung Ok Kim, Kyle Vaughn Laster, Xin Li, Baoyin Yuan, Zigang Dong, Kangdong Liu

## Abstract

**Background:** Esophageal cancer has a global impact on human health due to its high incidence and mortality. Therefore, there is an urgent need to develop new drugs to treat or prevent the prominent pathological subtype of esophageal cancer, esophageal squamous cell carcinoma. Based upon a screening of drugs approved by the Food and Drug Administration, we discovered that Arbidol could effectively inhibit the proliferation of esophageal squamous cell carcinoma *in vitro* and *in vivo*.

**Method:** We first conducted a series of cell-based assays and found that Arbidol treatment inhibited the proliferation and colony formation ability of ESCC cells and promoted G1 phase cell cycle arrest. Phospho-proteomics experiments and *in vitro* kinase assays were subsequently performed using cell lysates derived from Arbidol-treated cells in order to identify the underlying growth inhibitory mechanism. Finally, a PDX model was used to verify the inhibitory effect of Arbidol *in vivo*.

**Finding:** Arbidol inhibits proliferation of ESCC *in vitro* and *in vivo* through the DNA replication pathway and is associated with the cell cycle.

**Interpretation:** Arbidol is a potential ATR inhibitor via binding to ATR kinase to reduce the phosphorylation and activation of MCM2 at Ser108.

**Funding:** This research was supported by the National Natural Science Foundation of China (grant number 81872335), the National Natural Science Youth Foundation (grant number 81902486), and the Natural Science Foundation of Henan (grant number 161100510300).

## 1. Introduction

Esophageal cancer is a malignant neoplasm with a high incidence worldwide, ranking eighth in global incidence and sixth in overall mortality [1],[2]. It has two pathological types: esophageal squamous cell carcinoma (ESCC) and esophageal adenocarcinoma (EAC) [3]. ESCC is the main histological type and accounts for approximately 90% of esophageal cancers in China [4]. Current treatment options for patients with ESCC are surgery, chemotherapy, and radiotherapy [5]. However, the 5-year survival rate remains less than 20% [6]. Therefore, there is an urgent need for effective therapeutic or chemopreventive agents to improve the survival rate of patients with ESCC.

Chemoprevention is a promising strategy to prevent the recurrence of cancer [7]. In recent years, studies have found that certain non-steroidal anti-inflammatory drugs can be used for this purpose [8]. For example, it was reported that aspirin was used for chemoprophylaxis of colorectal, prostate, and other cancers [9]. Aside from non-steroidal anti-inflammatory drugs, other repurposed drugs also exhibit anti-tumor effects. Metformin, a hypoglycemic drug, has been shown to have a preventive effect on lung cancer, gastric cancer, pancreatic cancer, breast cancer, ovarian cancer, colorectal cancer, liver cancer, and other cancers [10],[11]. Therefore, it is likely that repurposing pre-existing Food and Drug Administration (FDA)-approved drugs could also be a promising strategy to prevent ESCC recurrence.

We screened a series of FDA-approved drugs and found that Arbidol has a significant inhibitory effect on the proliferation of ESCC cells. Arbidol is a broad-spectrum antiviral compound which is conventionally used for the prevention and treatment of influenza [12]. Its antiviral activity against a variety of DNA and RNA viruses is mainly based on disrupting key steps during the virus-cell interaction [13]. However, its role in cancer prevention or treatment has not been reported. Here, we identified that Arbidol is an inhibitor of the ataxia telangiectasia and Rad3-related (ATR) protein kinase.

ATR belongs to the phosphoinositide (PI) 3-kinase-related protein kinase (PI3KKs) family and one of its key functions is to regulate DNA replication [14],[15]. ATR can phosphorylate the key replication protein MCM2 (Minichromosome maintenance protein 2) on serine 108 in response to various forms of DNA damage and stalling of replication forks [15],[16]. In recent years, ATR inhibitors have been used in many cancer investigations [17], and their role in cancer treatment has gradually been realized.

We performed mass spectrometry (MS) analysis of lysates derived from Arbidol-treated cells to investigate the proteomic and phospho-proteomic response upon treatment. Additionally, we performed *in silico* molecular docking on a panel of kinases to predict potential binding partners and clarify the drug’s inhibitory mechanism. Our results showed that Arbidol inhibited ESCC growth via modulating the MCM2-ATR axis.

## 2. Materials and Methods

### 2. 1 Reagents

Arbidol was obtained from Beijing Ruitaibio Technology Co, Ltd (AT3360). ATR protein was purchased from SignalChem (No. A27-35G). Anti-rabbit ATR antibody was purchased from Cell Signaling Technology (13934S). Anti-rabbit MCM2 (phosphoS108) antibody was purchased from Abcam (ab109271). Anti-rabbit MCM2 antibody was purchased from Abcam (ab108935).

### 2. 2 Cell Lines and Cell Culture

Shantou human embryonic esophageal (SHEE) cells were obtained from Shantou University. 293T cells and the KYSE150 and KYSE450 ESCC cell lines were obtained from Cell Bank of the Chinese Academy of Sciences (Shanghai, China). The KYSE150 cells were maintained in RPMI-1640 medium supplemented with 10% fetal bovine serum (FBS) and 1% penicillin/streptomycin. KYSE450 cells were maintained in RPMI-1640/DMEM supplemented with 10% FBS and 1% penicillin/streptomycin. The 293T cells were maintained in RPMI-1640/DMEM supplemented (10% FBS). All cell lines were authenticated and periodically monitored for mycoplasma contamination.

### 2. 3 Cell Toxicity Assay

KYSE150 (8.0×10^3^ cells/well) and KYSE450 (1.0×10^4^ cells/well) cells were seeded into a 96-well plate with 100 μL of complete growth medium (with 10% FBS) containing various concentrations of Arbidol (final concentration 0, 6.25, 12.5, 25, 50, and 100 μM) and incubated for 24 h and 48 h. The cells were fixed with 100 μL of 4% paraformaldehyde at room temperature for 30 min. Next, 100 μL of 4’,6-diamidino-2-phenylindole (DAPI) solution was added to each well and the cells were incubated in the dark at 37°C for 20 min. The DAPI solution was subsequently discarded and the cells were washed twice with 1× phosphate buffered saline (PBS). Each well was then filled with 100 μL of fresh 1× PBS and high-content cell imaging analysis was used to photograph and analyze the number of cells. [18]

### 2. 4 Cell Proliferation Assay

KYSE150 (3.0×10^3^ cells/well) and KYSE450 (4.0×10^3^ cells/well) cells were seeded into a 96-well plate with 100 μL of complete growth medium (10% FBS) containing various concentrations of Arbidol (final concentration 0, 2.5, 5, 10, and 20 μM) and incubated at 37°C and 5% CO_2_ for 24 h, 48 h, 72 h and 96h. The cells were fixed with 4% paraformaldehyde at room temperature for 30 min. Next, 100 μL of DAPI solution was added to each well and the cells were incubated in the dark at 37°C for 20 min. The DAPI working solution was subsequently discarded and the cells were washed twice with 1× PBS. Each well was then filled 100 μL of fresh 1× PBS and high-content cell imaging analysis was used to photograph and analyze the number of cells. [18]

### 2. 5 Soft Agar Assay

Basal Medium Eagle (Sigma-Aldrich, UK) supplemented with 10% FBS, 0.5% agarose, 9% sterile water, 1% glutamine, 0.1% gentamicin, and various concentrations of Arbidol hydrochloride (final concentration 0, 2.5, 5, 10, and 20 μM) was added to each well of a 6-well plate, at 3 mL per well. After 2 h, cells (8×10^3^ per well) were suspended in 10% FBS, 0.33% agar, 45% sterile water, 1% glutamine, 0.1% gentamicin, and various concentrations of Arbidol (0, 2.5, 5, 10, and 20 µM). The cell suspension mixture was then applied over the solidified bottom layer. The cells were incubated at 37°C and 5% CO_2_ for 8 days. Afterward, the number of colonies was counted using an In Cell 6,000 analyzer.

### 2. 6 Colony Formation Assay

Cells were plated into 6-well plates at 200 cells per well and allowed to adhere for 12-16 h. Upon adherence, the cells were treated with various concentrations of Arbidol (0, 2.5, 5, 10, and 20 µM) for 10-14 days. The cell colonies were fixed with 4% triformol, stained with 0.1% Crystal Violet for 5 min, and then the number of colonies were counted.

### 2. 7 Western Blot Analysis

Cells were treated with various concentrations Arbidol for 24 h. The culture medium was then discarded and the cells were washed with 1x PBS prior to collection in radio immunoprecipitation buffer supplemented with phenylmethylsulfonyl fluoride and phosphatase inhibitor cocktail. Cell lysates were incubated for 1 h on ice, sonicated, and centrifuged for 30 min at 14,000 rpm and 4°C. Protein concentration was determined using the bicinchoninate method. The protein sample was resolved via sodium dodecyl sulfate polyacrylamide gel electrophoresis and transferred onto polyvinylidene difluoride membranes in transfer buffer. The membranes were incubated with the indicated antibodies prior to detection using enhanced chemiluminescent reagent.

### 2. 8 Pull Down Assay

Activated beads were suspended with 1 mM HCl, and a 40% dimethyl sulfoxide/H_2_ O (v/v)-coupling buffer [(pH 8.3) and 0.5 M NaCl] and then mixed with Arbidol (4 mg) prior to rotation overnight at 4°C. The beads were transferred to 0.1 M Tris-HCl buffer (pH 8.0) and again rotated overnight at 4°C. Finally, the beads were washed three times with 0.1 M acetate buffer (pH 4.0) containing 0.5 M NaCl followed by one additional wash with 0.1 M Tris-HCl (pH 8.0) containing 0.5 M NaCl [19].

### 2. 9 Molecular docking assay

The 2D structure of Arbidol (PubChem CID: 131411) was downloaded from the PubChem website (https://pubchem.ncbi.nlm.nih.gov/).. The crystal structure of ATR (5X6O) was downloaded from the Protein Database (PDB) website (https://www.rcsb.org/). The water molecules and small molecule ligands in the structure were deleted prior to the addition of hydrogen atoms. Computational docking between Arbidol and ATR was performed using the AutoDock 4.2.6 software. The optimal complex was visualized using PyMOL (PyMOL molecular graphics system, version 2.3.4).

### 2. 10 In vitro Kinase Assay

Purified MCM2 protein was used as the substrate for the *in vitro* kinase assay, with 200 ng active ATR (Signalchem). Reactions were performed in 1× kinase buffer (25 mM Tris-HCl pH 7.5, 5 mM β-glycerophosphate, 2 mM DTT, 0.1 mM Na_3_VO_4_, 10 mM MgCl_2_, and 5 mM MnCl_2_) containing 200 μM ATP at 30°C for 30 min. The reaction was terminated by adding 6 × protein loading buffer. The protein was detected using Western blotting.

### 2.11 Plasmids, Transfection, and Infection

ATR-specific short hairpin RNAs (shRNAs) were purchased from Sangon Biotech and then cloned into the pLKO.1 vector. The sequences were as follows: shATR#1: 5-GCCGCTAATCTTCTAACATTA shATR#2: 5-TAATGGTTCTTACTCGTATTA. To prepare ATR virus particles, we used jetPRIME to transfect each viral vector and packaging vector into HEK293T cells according to the protocol recommended by the manufacturer. After 48 h of incubation, the viral medium was collected and filtered through a 0.45 μm sodium acetate filter. The viral medium was then supplemented with 8 μg/mL polyethylene and used to infect KYSE150 or KYSE450 cells. After 24 h of incubation, cells were selected with puromycin (1 μg/mL) for 48 h.

### 2.12 PDX Mouse Model

6-8 week old severe combined immunodeficiency (SCID) female mice (Vital River Labs, Beijing, China) were used for patient-derived xenograft (PDX) mouse model experiments in accordance with the guidelines of the Ethics Committee of Zhengzhou University (Zhengzhou, Henan, China),. The EG 20 ESCC case was used for the PDX animal experiments. The tumor was divided into 0.1-0.2 g fragments and inoculated onto the backs of SCID mice. The mice were randomly divided into the following groups: control group, 100 mg/kg Arbidol hydrochloride treatment group, and 200-400 mg/kg Arbidol hydrochloride treatment group. Arbidol hydrochloride or dimethyl sulfoxide was administered to each mouse daily by oral gavage. The tumor volume and weight of each mouse were measured every 3 days. The tumor volume was calculated according to the size of a single tumor and the following formula: tumor volume (mm^3^) = 1/2 × length × width^2^. [20]

### 2.13 Cell Cycle Assay

KYSE150 cells and KYSE450 were cultured in 6-well plates at a density of 8×10^5^ and 1.2×10^6^ cells/well, respectively. After serum starvation for 24 h, treated with Arbidol. The cells were then harvested and fixed in 70% cold ethanol at 4°C overnight. After fixation, the cells were washed three times with PBS, stained with propidium iodide staining solution for approximately 30 min at room temperature, and analyzed using FACScan flow cytometer. Data was analyzed using FlowJo.

### 2.14 Liquid Chromatography Tandem Mass Spectrometry (LC-MS/MS) Analysis

The peptide was dissolved in solvent A (0.1% formic acid and 2% acetonitrile in water) and separated using an ultra-high-performance liquid chromatography (UPLC) system (EASY-nLC 1000; Thermo Scientific). The gradient included an increase from 6% to 22% solvent B (0.1% formic acid in 90% acetonitrile) for 40 min, 22% to 35% solvent B for 12 min, 35% to 80% solvent B for 4 min, and rising to 80% solvent B for the last 4 min. A constant flow rate of 700 nL/min was used. The separated peptides were then ionized using a nanospray ionization ion source and analyzed using an Orbitrap Fusion™ (Thermo Scientific) mass spectrometer. High-resolution Orbitrap was used to detect and analyze the precursor ions and secondary fragments of peptides. The m/z scan range of the full scan was 350-1550, and the complete peptide was detected in the Orbitrap at a resolution of 60,000. The scanning range of the secondary mass spectrometer was fixed at the starting point of 100 m/z, and the resolution was set to 15,000. After a full scan, the top 20 peptides with the highest signal intensity were selected to enter the high-energy collision dissociation library and fragmented using 35% of the fragmentation energy. A secondary mass spectrometry was also performed.

### 2.15 Gene Set Enrichment Analysis (GSEA) For Data Processing

The GSEA v4.0.3 software package was used to integrate the proteome data set with gene set enrichment analysis (GSEA) and to determine the difference in protein expression between the treatment and control groups. The mitochondrial protein complex genome (GO: 0098798) and the respiratory electron transport genome (R-HSA-611105) were selected for enrichment. Through previous functional annotations or experiments, GSEA determined the relevant gene set and sequenced the genes based on the degree of differential expression of the two samples. Gene set enrichment analysis detected changes in gene set expression and optimized the results using P value and false discovery rate (FDR) methods. Biological pathways with P <0.01 and FDR <0.25 were considered statistically significant.

### 2.16 Immunohistochemical (IHC) Analysis

The IHC assay was carried out as described previously [20]. Briefly, paraffin-embedded tissues (5 μm) were prepared for IHC analysis. The specimens were baked in a constant-temperature oven at 65°C for 2 h, de-paraffinized, and rehydrated prior to antigen retrieval. All samples were treated with 3% H_2_O_2_ for 10 min in the dark to inactivate endogenous peroxidases. The slides were then incubated with specific antibodies overnight at 4°C. The tissue sections were washed three times with 1× Tris-buffered saline with 0.1% Tween ® 20 Detergent and hybridized with the secondary antibody for 15 min at 37°C. After DAB staining, all slides were stained with hematoxylin, dehydrated, and mounted under glass coverslips.

### 2.17 Statistical Analysis

All quantitative results are expressed as mean ± SD. Statistically significant differences were checked using Student’s t-test or one-way analysis of variance. p<0.05 was considered statistically significant. \**P* < 0.05, \*\**P* < 0.01, \*\*\**P* < 0.001.

## 3. Results

### 3.1 Arbidol inhibits ESCC cell proliferation

To identify a drug able to inhibit ESCC cell proliferation, we conducted toxicity experiments using FDA-approved drug libraries and discovered that Arbidol (Figure 1a) effectively suppresses ESCC cell growth (Figure 1b). The half-maximal inhibitory concentration (IC50) concentrations of the drug in KYSE150, KYSE450, and SHEE cells were 57.54, 54.72, and 87.63 μM at 24 h, and 26.37, 20.75, and 62.52 μM at 48 h, respectively (Figure 1c). To further evaluate the effect of Arbidol on ESCC growth, we treated KYSE150 and KYSE450 cells with different concentrations (0, 2.5, 5, 10, and 20 µM) of Arbidol (Figure 1d). The results indicated that Arbidol inhibited the proliferation of KYSE150 and KYSE450 cells in a dose-dependent manner. It is worth mentioning that we also performed toxicity and proliferation experiments on the human immortalized SHEE esophageal epithelial cell line and observed that the growth inhibitory effect was less pronounced. We next assessed whether treatment with Arbidol could impact anchorage-independent cell growth using the soft agar assay. The result showed that Arbidol inhibited anchorage-independent growth of KYSE150 and KYSE450 cells in a dose-dependent manner (Figure 1e). Finally, we used the colony-formation assay to determine whether Arbidol inhibited the proliferation of ESCC cell lines (Figure 1f). Similarly, the results indicate that Arbidol inhibited colony-formation in a dose-dependent manner. Collectively, these results suggest that Arbidol has a significantly inhibitory effect on ESCC *in vitro*.

**Figure 1.**
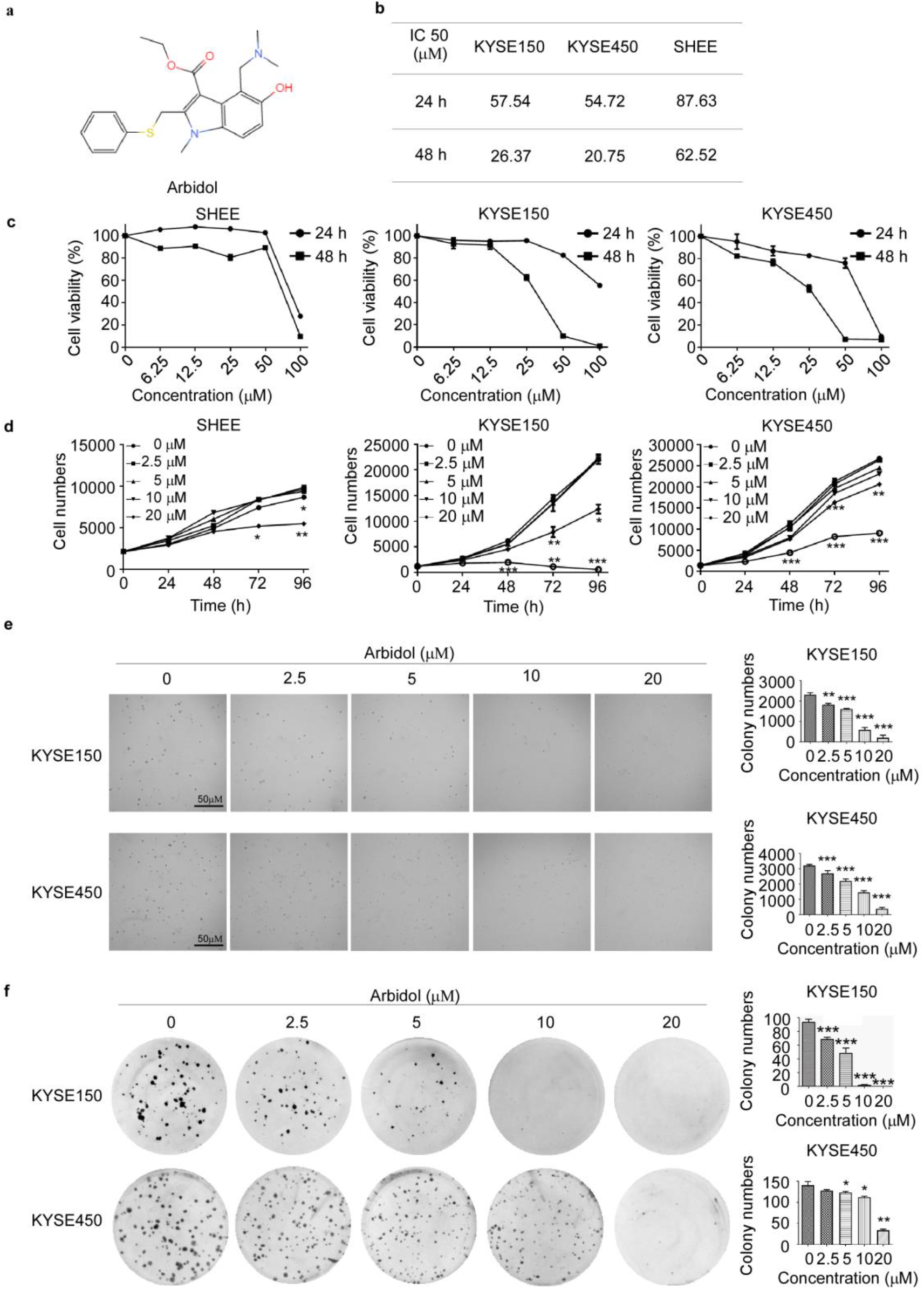
Arbidol inhibits ESCC cell proliferation. a. The structure of Arbidol. b. KYSE150, KYSE450, and SHEE cell viability was detected by IN Cell Analyzer. KYSE150 and KYSE450 cells were treated with different concentrations of Arbidol (0, 6.25, 12.5, 25, 50, and 100 μM) for 24 h and 48 h and evaluated by 4′,6-diamidino-2-phenylindole. IC50 was calculated using SPSS. c. KYSE150, KYSE450, and SHEE cells were treated with Arbidol (0, 6.25, 12.5, 25, 50, and 100 μM) at different concentrations as indicated for 24 h and 48 h. Cell viability was then determined by IN Cell Analyzer. d. The effect of different concentrations of Arbidol (0, 2.5, 5, 10, and 20 μM) on SHEE, KYSE150, and KYSE450 cells for 0, 24, 48, 72, and 96 h. Asterisks (*, **, ***) indicate a significant decrease (*P < 0.05, **P < 0.01, ***P < 0.001, respectively). e. Arbidol effectively inhibits anchorage-independent cell growth. Colony numbers were counted using an IN Cell Analyzer. Asterisks indicate a significant difference compared with the control group. f. Arbidol effectively inhibits colony formation in esophageal squamous cell carcinoma (ESCC) cell lines. KYSE150 (200/well) and KYSE450 (400/well) cells were seeded in 6-well plates and treated with different concentrations of Arbidol (0, 2.5, 5, 10, and 20 μM). The numbers of cell colonies were counted after a period of time. Asterisks indicate a significant difference (*P <0.05, ** P <0.01, and ***P <0.001) compared with the control group.

### 3.2 Phosphoproteomics reveals that Arbidol inhibits tumors through the MCM-ATR signaling pathway

To determine the specific inhibitory mechanism of Arbidol in ESCC, KYSE150 cells were treated with 20 μM arbidol or dimethyl sulfoxide as control for 24 h. Subsequently, mass spectrometry was performed (Figure 2a). We identified 241 down-regulated proteins and 340 down-regulated sites. Next, we performed KEGG enrichment analysis using the quantified phosphate sites to query significantly down-regulated pathways (Figure 2b, 2c). DNA replication was identified as the key pathway affected. A heat map detailing phosphorylation site enrichment and KEGG pathway enrichment showed that the S108 site of MCM2 in the DNA replication pathway was significantly down-regulated after Arbidol treatment (Figure 2d, 2e). The results of the *in silico* were verified by Western blotting (Figure 2f). The Western blotting results confirmed the phospho-proteomics data and showed that MCM2 S108 phosphorylation was strongly inhibited after Arbidol treatment. The peptide figure showed that the test conformed the standard (Figure 2g). To further investigate the specific targets of Arbidol, we first searched for proteins upstream of P-MCM2 Ser108 and found that Cdc7 and ATR can phosphorylate MCM2 (Ser108) [21] [22] [23]. However, pull-down results indicated that Cdc7 could not directly bind to Arbidol (Supplementary a). Next, we conducted an in-depth analysis of the recent quantitative human interaction group (pull-down mass spectrometry), and utilized the kinase prediction tool (from Phosphoproteomics data). We also used the chemical structure of Arbidol to predict targets using the web tool Swiss Target Prediction. The results suggested that ATR kinase was a potential target of Arbidol.

**Figure 2.**
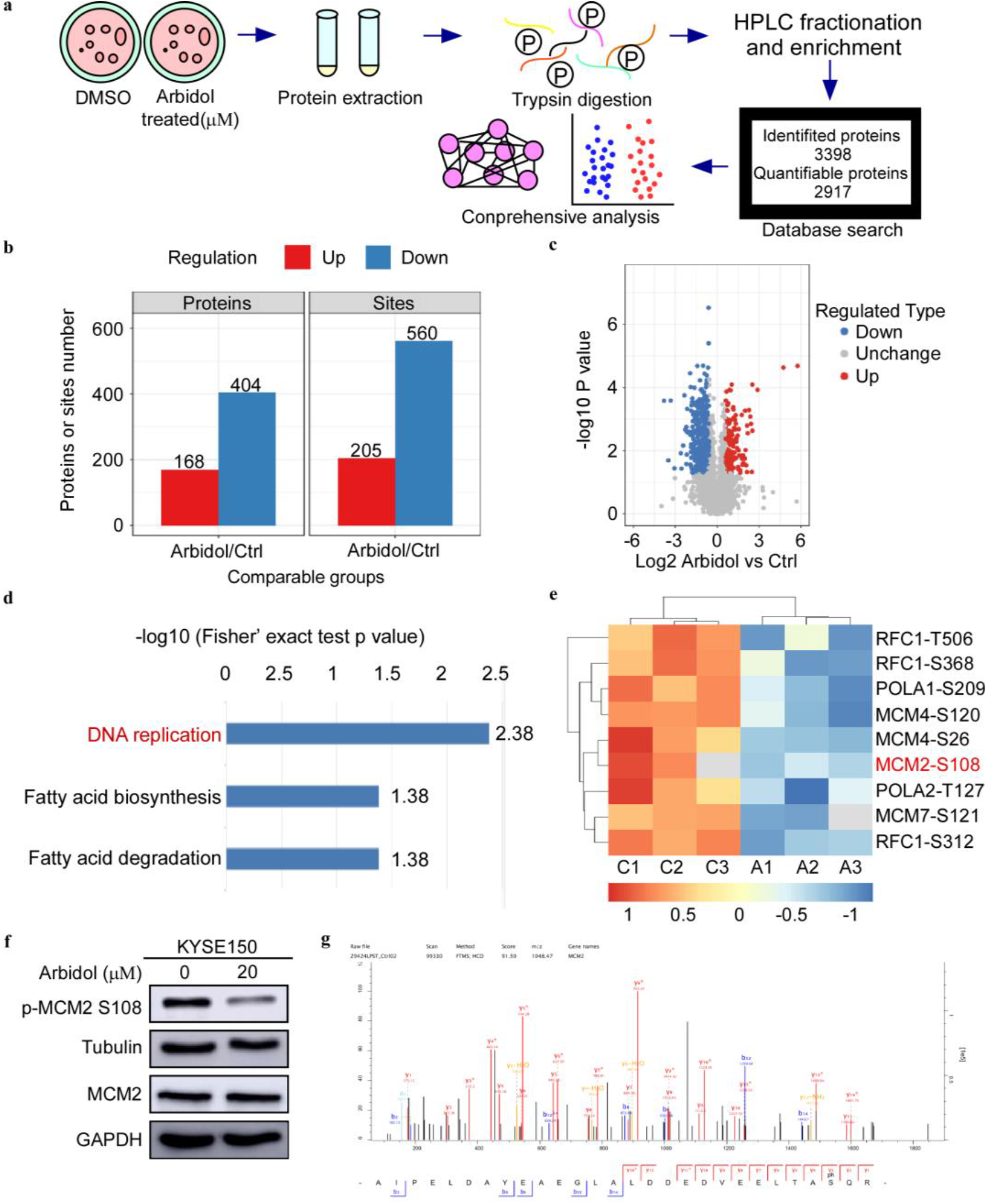
Phosphoproteomics reveals that arbidol inhibits tumors through minichromosome maintenance-ataxia telangiectasia and Rad3-related (MCM-ATR) protein. a. The mass spectrometry workflow of phosphorylated proteomics analysis of KYSE150 cells after 24 h treatment with Arbidol. b. The histogram shows the regulation of phosphorylation sites compared to the control group. c. The volcano plot shows that 375 phosphorylation sites changed significantly. d. Kyoto Encyclopedia of Genes and Genome (KEGG) analysis indicated that three down-regulated pathways were enriched. e. Phosphorylation proteomics analysis of the esophageal squamous cell carcinoma (ESCC) cell line after 20 μM Arbidol treatment. The different phosphorylation sites are plotted as heat maps. f. Western blot analysis shows that the phosphorylation of MCM2 S108 is markedly downregulated by Arbidol. g. Distribution of peptide mass error and length of protein peptides identified in this study.

### 3.3 Arbidol targets ATR to affect DNA replication pathway in ESCC

We conducted a computational docking study using the molecular structures of Arbidol and ATR to determine optimal binding orientation. The results suggested that Arbidol could bind to ATR at the ASN 2346 and ASN 2361 residues (Figure 3a). Based on the sample information from pull-down mass spectrometry, we found that 906 candidate targets were differentially represented between the control and treatment groups. Next, we verified that one of the targets of ATR could bind to Arbidol through a pull-down assay *in vitro* and *ex vivo* (Figure 3b-3d). To validate the results of the molecular docking as well as to assess the contribution of ASN 2346 and ASN 2361 to the binding between Arbidol and ATR, we mutated these two residues and performed a pull-down assay. The results showed that the binding between Arbidol and ATR was less efficient after the mutation (Figure 3e). In addition, the results of an *in vitro* kinase assay indicated that Arbidol treatment reduced the phosphorylation of MCM2 (Figure 3f). The ATP competition assay further illustrated that Arbidol functions as an ATP-competitive inhibitor of ATR (Figure 3g). These results indicated that ATR was involved in the phosphorylation of MCM2, and that ATR inhibition induced by Arbidol treatment may affect the phosphorylation of MCM2.

**Figure 3.**
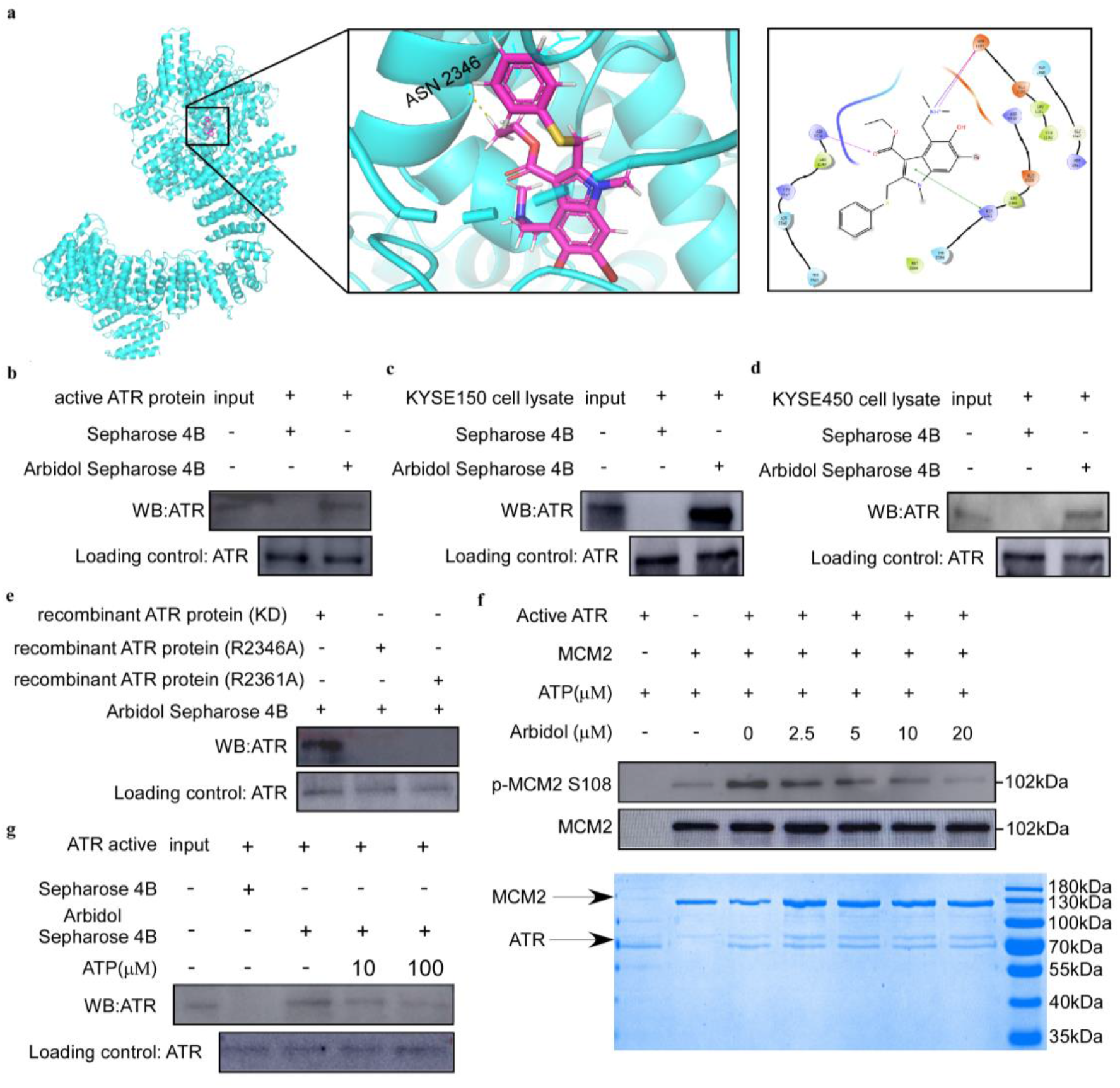
Arbidol targets ataxia telangiectasia and Rad3-related (ATR) protein to affect the DNA replication pathway in ESCC. a. Computational docking was used to identify the binding site of Arbidol and ATR. b-d. Arbidol directly binds to ATR. The ESCC cell lysate (500 μg) (c, d) and ATR pure protein (1 μg) (b) were incubated with sepharose 4B beads conjugated with Arbidol or sepharose 4 B beads alone. Western blot analysis was used to analyze the pull-down protein. e. The Arbidol pull-down assay and amino acid site mutation of ATR protein. f. *In vitro* kinase assay. Arbidol attenuates the phosphorylation of minichromosome maintenance 2 (MCM2) by inhibiting the kinase activity of ATR. g. ATP competition assay. ATR pure protein (1 μg) was incubated with Sepharose 4 B beads conjugated with Arbidol, and with ATP (10 μg or 100 μg) or Sepharose 4 B beads alone. Western blot analysis was used to analyze the pull-down protein.

### 3.4 Arbidol inhibits the proliferation of ESCC cells and arrests the cell cycle at G1 phase

To elucidate the functional pathways of ATR, a GSEA analysis was performed using our phosphoproteomics and proteomics datasets (Figure 4a). We identified MCM2 downstream of ATR in the cell cycle pathway, thereby confirming a relationship between them. We then performed Western blotting and verified that the phosphorylation of MCM2 S108 decreased upon treatment with Arbidol in a dose-dependent manner (Figure 4b). Variations in DNA replication timing are often accompanied by changes in the cell cycle. We used propidium iodide staining to examine the effect of Arbidol on cell cycle progression in KYSE150 and KYSE450 cells after treatment for 48 h. Results showed that Arbidol strongly reduced the S phase fraction and induced cell cycle arrest at the G1 phase in a dose-dependent manner (Figure 4c).

**Figure 4.**
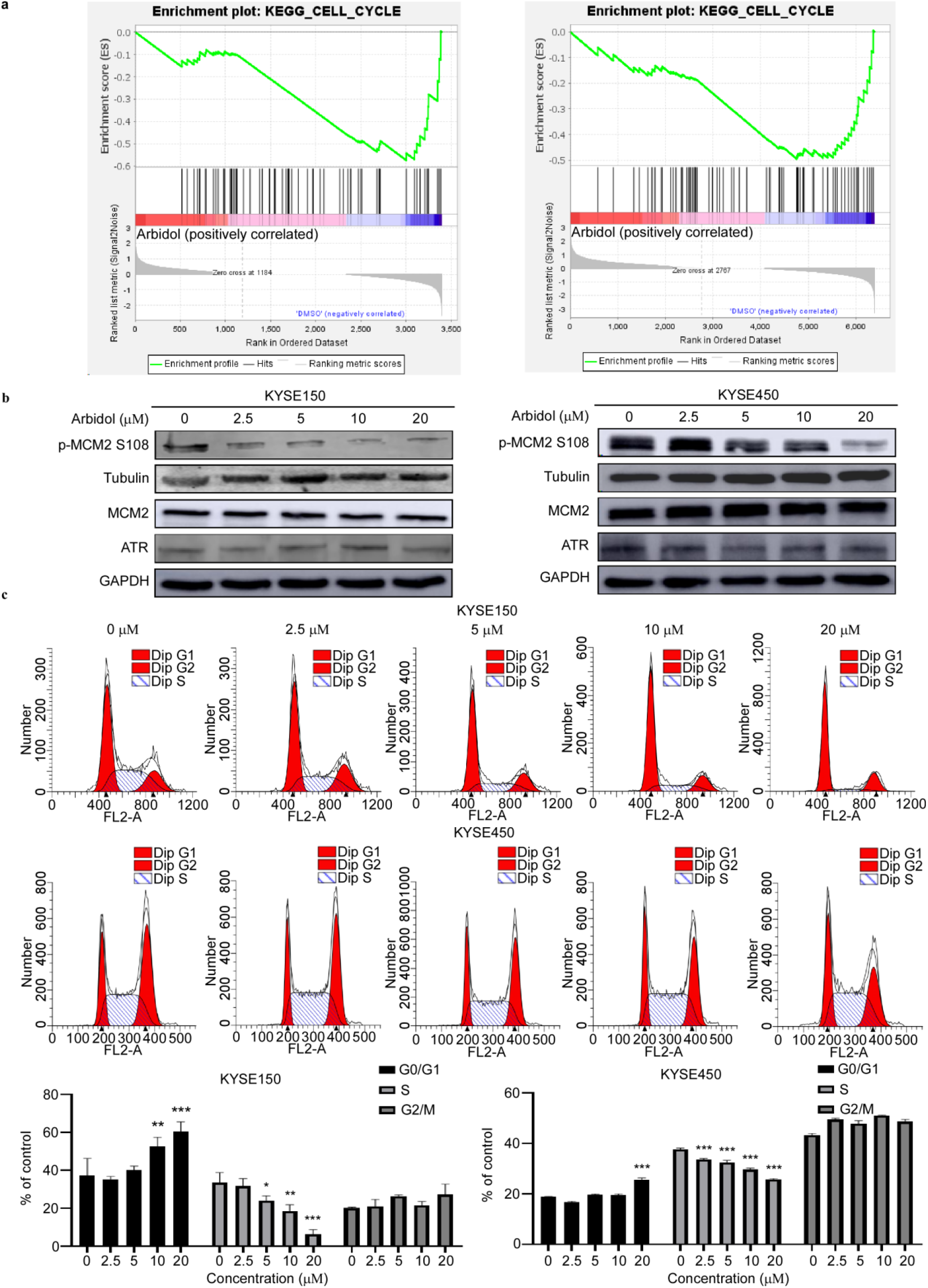
Arbidol inhibits the proliferation of ESCC cells and arrests the cell cycle in G1 phase. a. Gene set enrichment analysis (GSEA) indicates the representation of the cell cycle pathway in both the phosphoproteomics and proteomics datasets. b. The expression level of p-MCM2 S108 decreases with increasing concentrations of Arbidol in KYSE150 and KYSE450 cells (0, 2.5, 5, 10, and 20 μM), as confirmed by western blotting. c. After 24 h of Arbidol treatment, KYSE150 and KYSE450 cells were stained with propidium iodide (PI) and the cell cycle distribution was analyzed using flow cytometry. The quantitative cell cycle distribution data of KYSE150 cells are shown here.

### 3.5 ATR knockdown reduces the growth of ESCC cells

Next, we determined the correlation between ATR and MCM2 expression using the Bioanalysis website (https://www.aclbi.com/static/index.html#/). The results showed a high correlation between ATR and MCM2 expression (Figure 5a). We next queried the TCGA (The Cancer Genome Atlas) database to analyze the expression profile of ATR in esophageal tumor and normal tissues based on sample types or tumor history (Figure 5b). We also employed timer 2.0to analyze the expression status of ATR across tissue samples. (Figure 5c). These results support that ATR high expression correlates with decreased overall survival. To determine the effect of ATR knockdown on the growth of esophageal cancer cells (KYSE150 and KYSE450), we established ATR stable knockdown cells and confirmed the expression of ATR protein by western blotting. The results showed that the expression of total ATR in sh*ATR*#3 and sh*ATR*#5 cells was greatly reduced (Figure 5d). We then examined the effect of ATR knockdown (KD) on the growth of esophageal cancer cells using a proliferation assay. The results showed that ATR KD significantly reduced the growth of esophageal cancer cells (Figure 5e, 5f). To determine whether the growth reduction of esophageal cancer cells observed upon treatment of Arbidol depended on the expression of ATR, we performed a proliferation assay using ATR KD cells treated with various concentrations (0, 2.5, 5, 10, and 20 µM) of Arbidol. The results showed that cells expressing sh-ATR were resistant to the growth inhibitory effects of Arbidol compared with cells expressing sh-MOCK (Figure 5g).

**Figure 5.**
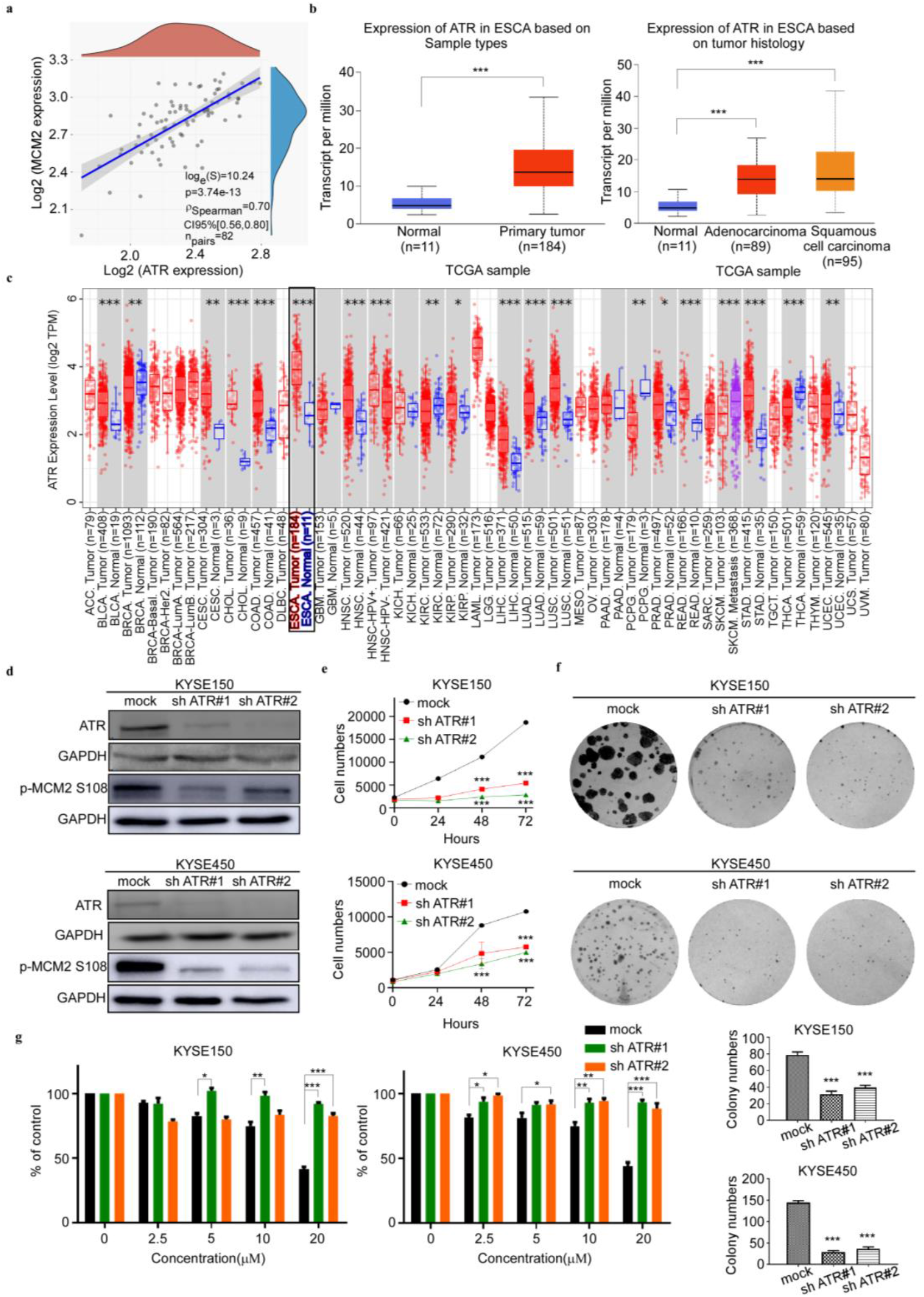
Knocking down of ataxia telangiectasia and Rad3-related (ATR) reduces the growth of ESCC cells. a. The correlation between ATR and minichromosome maintenance 2 (MCM2) is detected by gene correlation analysis. b. TCGA data results showing the expression of ATR in different cancer types, including esophageal cancer and normal tissue, as well as the statistical chart illustrating esophageal cancer classification. c. The picture is from the results of bioassay on the association between ATR and pan-cancer on Timer 2.0, which includes esophageal cancer. d. KYSE150 and KYSE450 cell lines stably expressing knockdown sh-*ATR* or shControl were established. The expression of ATR and MCM2 S108 is confirmed by western blotting. e. The effect of ATR gene knockdown on the growth of esophageal cancer cells. The stable knockdown sh-*ATR* cells were seeded into 96-well plates, fixed at 0 h, 24 h, 48 h, 72 h, and 96 h, stained with 4′,6-diamidino-2-phenylindole, and analyzed by IN Cell Analyzer. f. The effect of ATR gene knockdown on the clonogenic potential of esophageal cancer cells. KYSE150 (200/well) and KYSE450 (400/well) cells were seeded on a 6-well plate. The number of cell colonies were counted after 15 days. Asterisks indicate a significant difference (*P <0.05, P <0.01, and ***P <0.001) compared with the control group. g. The effect of Arbidol on the growth of ESCC cells was evaluated in cells stably expressing sh-*ATR* or cells stably expressing MOCK. After inoculating the cells for 12 h, they were in the presence or absence of different concentrations of Arbidol, and analyzed with IN Cell Analyzer 96 h later.

### 3.6 Arbidol inhibits tumor growth of ESCC patient-derived xenograft in vivo

To test whether Arbidol exerts an inhibitory effect on ESCC cells *in vivo*, we utilized an ESCC PDX mouse model (Figure 6a). The results of the experiment showed that Arbidol treatment effectively reduced the tumor volume and weight of EG 20 xenografts (control group, n = 7; low Arbidol group, n = 7; high Arbidol group, n = 7; Figure 6b). Importantly, no significant difference in body weight was observed between the control and experimental groups. (Figure 6c-6f). Compared with the control group, histological staining confirmed that Arbidol substantially impaired MCM2 S108 expression in the case of EG 20 models, and reduced the expression levels of cell proliferation marker Ki67 (Figure 6g).

**Figure 6.**
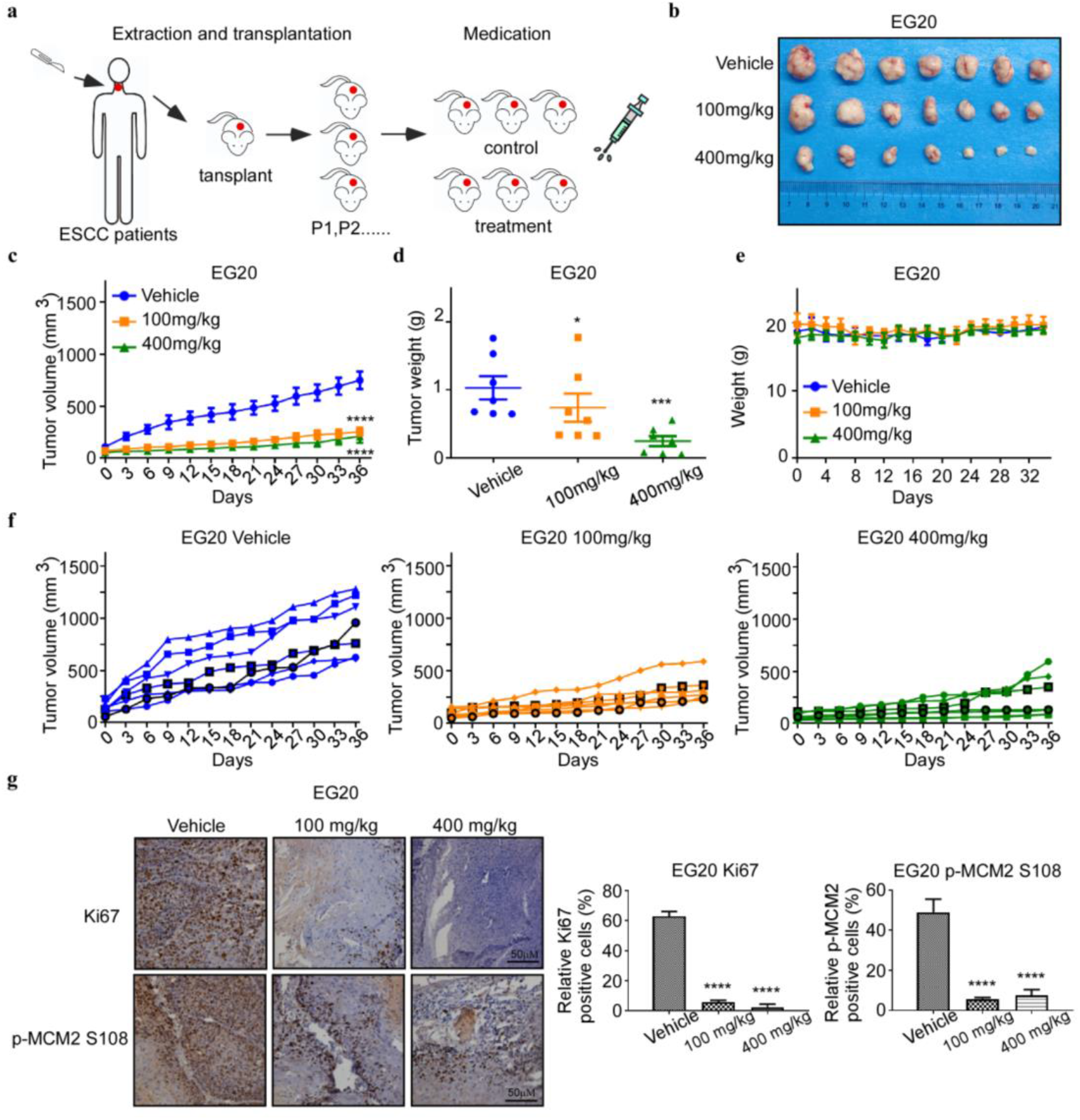
Antitumor efficacy of Arbidol in an ESCC patient-derived xenograft model. a. Arbidol treatment protocol for ESCC PDX models. b. Tumor sizes of the EG 20 xenografts are shown. Tumors were excised and weighed at the end of the experiment (36 days after treatment). c. The effect of Arbidol inhibition on *in vivo* tumor growth was determined using ESCC patient-derived xenograft models: EG 20. Mice (n = 7 per group) were treated with vehicle (saline, oral administration, daily) or Arbidol (low = 100 mg/kg, high = 400 mg/kg, daily) for 36 days. Tumor volume was measured every 3 days. p values were calculated using a standard one-way ANOVA. Data are presented as the mean. d. Tumor growth curve of single mouse grafted with EG 20 is shown. n = 7 per group. e. Weight of mice treated with Arbidol. f. Tumor volume of each group was shown respectively. n = 7 per group. g. The expression of tumor proliferation markers Ki67 and target engagement was verified by immunohistochemical analysis of MCM2 S108 expression in Arbidol-treated EG 20 PDX mice. n = 7 per group.

## 4. Discussion

The five-year survival rate of ESCC is less than 20% due to limited preventive methods after primary treatment. Chemoprevention is a promising strategy to reduce the recurrence of ESCC. Several NSAIDs and anti-metabolites drugs have been shown to influence the recurrence rate of ESCC. However, few drugs have been successfully repurposed to clinically treat cancer [24]. In the present manuscript, we found that Arbidol significantly reduced ESCC cell proliferation, anchorage-dependent growth, and anchorage-independent growth *in vitro* (Figure 1d, 1e, 1f). Additionally, we used a PDX mouse model to evaluate the inhibitory effect of Arbidol *in vivo*. Our results indicated that Arbidol treatment resulted in a significant reduction of tumor size and weight compared to the vehicle-treated group. Importantly, no significant difference in body weight was observed between the treatment and control groups, suggesting that Arbidol could be considered a promising chemoprevention drug against ESCC.

ATR is a key component of the targeted DNA damage response (DDR) [25], a new cancer treatment pathway which has broad prospects for tumor selectivity [26]. ATR can phosphorylate proteins involved in DNA replication, DNA recombination and repair, and cell cycle regulation in response to DNA damage [27]. Due to its extensive role in DDR, ATR inhibition has great potential in cancer treatment [28],[29],[30]. Therefore, suppressing ATR signal transduction by inhibitors may exert anti-tumor effects against ESCC [31].

DNA replication occurs in all organisms that use DNA as their genetic material and their molecular basis of biological inheritance [32]. During DNA replication, a key component called the MCM complex binds to related proteins and provides the helicase activity that unwinds the DNA at the origin of replication [33],[34]. In the present study, we identified that ATR is an upstream kinase of MCM2 using a predictive model based on multiple proteomic assays. ATR was found to be a target of Arbidol through pull-down and *in vitro* kinase assays (Figure 3b-3f). We determined that Arbidol bound to ATR at Asn2346 and Asn2361 sites (Figure 3a), and that these residues are functionally important binding sites for the Arbidol and ATR interaction (Figure 3e). Additionally, these residues down-regulated the phosphorylation level of MCM2 (Figure 4b) and affected ATR kinase activity, as well as the cell cycle (Figure 4c).

According to our research, we found that Arbidol is a potential ATR inhibitor that effectively inhibits ESCC growth *in vitro* and *in vivo*. At present, the anti-tumor activity of ATR inhibitors has been observed in many preclinical studies, and the encouraging results of these studies have prompted clinicians to evaluate their effectiveness and safety in clinical trials [35]. Existing ATR inhibitors, such as AZD6738, VX-970/M6620, and BAY-1895344 are currently being tested in a series of clinical trials. Of the listed inhibitors, VX-970 is currently being investigated in a Phase I dose-escalation safety study for use to treat esophageal cancer and other solid cancers in combination with radiotherapy and chemotherapy [36]. However, the development of effective and specific drugs that target ATR is proceeding slowly [37]. Compared with the development strategy of de novo synthesis, FDA-approved drugs usually provide pharmacokinetic and toxicity data and are more convenient for clinical translation.

In summary, our research showed that Arbidol targets ATR, thereby attenuating ESCC proliferation *in vitro* and *in vivo*.

## Acknowledgments

Thanks to Xiaofan Zhang, Xiangyu Wu, Zhuo Bao, Bo Li and Yubing Zhou from Pathophysiology Department, Zhengzhou university.

## Authors’ contributions

Zigang Dong and Kangdong Liu developed the idea, Ning Yang participated in all experiments and wrote the manuscript, Yanan Jiang and Kyle Vaughn Laster revised the paper, Xuebo Lu and Donghao Wang performed the animal experiments, Lili Zhao performed the immunohistochemical staining experiment, Yaxing Wei conducted cell experiments, Yin Yu performed the protein purification work, Xin Li and Baoyin Yuan analyzed the data.

## Supplemental Material

**Fig. S1.**
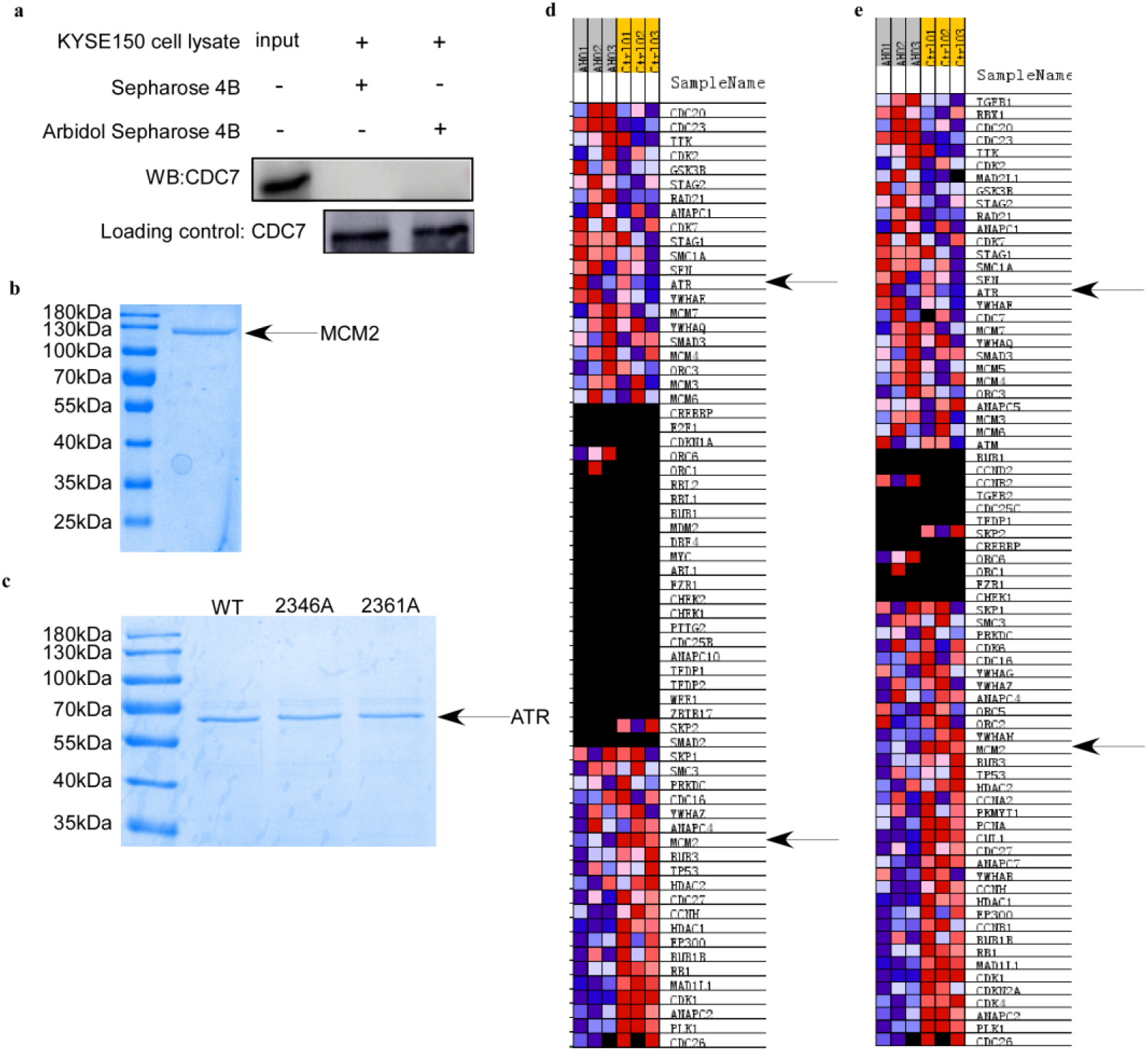
a. Arbidol does not bind directly to CDC7. The KYSE150 cell lysate (500 μg) was incubated with sepharose 4B beads conjugated with Arbidol or sepharose 4B beads alone. Western blot analysis was used to analyze the pull-down protein. b. Coomassie blue-stained gel of purified MCM2, the black arrow indicates MCM2 c. Mutant protein ATR purified by Coomassie Bright Blue staining gel, the first lane is wild type protein, the second and third lanes show mutant ATR protein. d, e. Gene set enrichment analysis (GSEA) includes the phosphoproteomics (left panel) and proteomics (right panel) datasets. The black arrow indicates MCM2 and ATR.

